# Ethylene-Gibberellin Crosstalk Drives Phenotypic Sex Changes in *Cannabis sativa*

**DOI:** 10.64898/2026.05.11.724340

**Authors:** Julien Roy, Adrian S. Monthony, Davoud Torkamaneh

## Abstract

Sex expression in *Cannabis sativa* is determined by XX/XY sex chromosomes but remains plastic, with ethylene inhibition inducing male flowers on XX plants and ethylene release inducing female flowers on XY plants. Although ethylene is a central regulator of this process, the contribution of gibberellin signaling to cannabis sex reversal remains poorly defined. Here, we reconstructed the GA biosynthesis, regulation, and signaling pathway in *C. sativa* and profiled GA-related gene expression during chemically induced sex reversal. Orthology-based searches identified 50 putative *C. sativa* GA-related genes, widely distributed across the genome, with the X chromosome harboring 11 genes, including six within the non-recombining region. Transcriptomic analyses across vegetative baseline, early post-treatment leaves, and developing flowers showed that expression profiles were broadly similar between XX and XY plants at day 0, weakly perturbed at day 1, and strongly structured by floral phenotype at day 14. Early responses were limited to downregulation of *CsGA3ox1* in ethephon-treated XY plants and *CsGASA1* in STS-treated XX plants. By day 14, sex reversal was associated with differential expression of key genes, including *CsGA1*, multiple *GA20ox* orthologs, *CsGID1B, CsSLY2*, and several *GASA* genes, indicating broad remodeling of GA regulation. Our findings position the GA pathway as a downstream module of ethylene-driven sex reversal in *C. sativa*, with GA activity tracking floral sexual identity, extending the framework of sexual plasticity beyond ethylene, and identifying candidate genes for functional validation and the development of sex-stable cultivars.

## Introduction

Sexual reproduction in angiosperms encompasses a remarkable diversity of breeding systems. Most flowering plant species produce bisexual (hermaphroditic) flowers bearing both male and female organs, a condition that facilitates self-pollination and reproductive assurance (Barrett, 2002; Renner, 2014). However, a minority of angiosperms have evolved unisexual flowers, giving rise to monoecious species, in which separate male and female flowers develop on the same individual, and dioecious species, in which male and female flowers are borne on distinct individuals (Charlesworth, 2002). Dioecy has evolved independently hundreds of times across the angiosperm phylogeny and occurs in approximately 5–6% of flowering plant species (Renner, 2014). In dioecious lineages, sex is often determined by sex chromosomes, which may range from homomorphic pairs with limited differentiation to highly heteromorphic systems resembling those of animals (Ming et al., 2011). The evolution of sex chromosomes in plants is thought to proceed through the progressive suppression of recombination around a sex-determining locus, leading to the accumulation of sex-linked genes and, eventually, morphological differentiation of the sex chromosome pair (Bergero & Charlesworth, 2009).

Despite the presence of genetic sex-determining mechanisms, sexual expression in many plant species is not rigidly fixed. In both dioecious and monoecious plants, the phenotypic sex of flowers can be modulated by environmental conditions, developmental stage, and phytohormone signaling, a phenomenon broadly referred to as sexual lability or sexual plasticity (Cossard & Pannell, 2021; Käfer et al., 2022). Sexual plasticity has been documented across diverse taxa, including *Carica papaya* (papaya), *Spinacia oleracea* (spinach), *Mercurialis annua*, and *Amborella trichopoda* (Anger et al., 2017; Cossard & Pannell, 2021; Lin et al., 2016). The triggers that shift phenotypic sex or floral sex ratios vary widely and include natural population variability, reproductive pressure, abiotic or biotic stress, and targeted molecular interventions with plant growth regulators (Dennis Thomas, 2004; J. Zhang et al., 2017). Among the phytohormones implicated in sex determination, ethylene and gibberellins (GAs) have emerged as central players, with evidence from multiple plant families linking their biosynthesis and signaling to the control of male versus female flower development (Chailakhyan & Timiriazev, 1979; Chandler, 2011; Diggle et al., 2011).

*Cannabis sativa* L. is a predominantly dioecious species with male and female flowers on separate individuals (Bonini et al., 2018). The species has 10 chromosome pairs (2n = 20), comprising nine autosomal pairs and one sex-chromosome pair: males are typically XY with heteromorphic sex chromosomes, whereas females are XX with a homomorphic pair (Carey et al., 2024; Prentout et al., 2020). *C. sativa* develops sexually dimorphic flowers. Pistillate flowers comprise an ovary enclosed by two bracts bearing glandular trichomes and terminate in two elongated stigmas (Leme et al., 2020; Spitzer-Rimon et al., 2019). Staminate flowers consist of a simple perianth and typically five stamens positioned opposite the sepals (Schilling et al., 2020). The glandular trichomes found in female inflorescences produce a diversity of cannabinoids with substantial pharmacological interest (Andre et al., 2016). In recent years, the cannabis genome has been sequenced and assembled (Grassa et al., 2021; Van Bakel et al., 2011), and interest in the genetic and molecular basis of sex determination has grown considerably (Adal et al., 2021; Chen et al., 2025; Monthony et al., 2026; Orozco et al., 2025; Prentout et al., 2020; Shi et al., 2025).

As observed in other dioecious plants, sex expression in *C. sativa* is not determined solely by sex chromosomes. Under specific environmental or experimental conditions, cannabis plants can display flowers that do not align with their genotypic sex, a phenomenon that has been recognized for decades (Chailakhyan & Timiriazev, 1979; Mohan Ram & Sett, 1982). Evidence for hormonal regulation of cannabis sexual plasticity comes from early studies using exogenous plant growth regulators, which demonstrated that the application of gibberellins, particularly GA_3_, to genetically female (XX) plants could induce the formation of male flowers (Galoch, 1978; Ram & Jaiswal, 1972). Conversely, ethylene emerged as a key modulator of feminization: inhibition of ethylene signaling using silver thiosulfate (STS) or silver nitrate reliably induces male flowers on genotypically female plants (induced male flowers; IMF), whereas exogenous applications of ethylene or the ethylene-releasing compound ethephon induce feminization of XY plants (induced female flowers; IFF) (Flajšman et al., 2021; Garcia-de Heer et al., 2025; Monthony et al., 2026; Moon et al., 2020a). These chemical interventions now form the basis of feminized seed production in the cannabis industry, yet the underlying molecular mechanisms governing chemically induced sex-change have remained poorly understood until recently.

Recent transcriptomic and multi-omics investigations have begun to uncover the molecular architecture of ethylene-mediated sexual plasticity in cannabis. A comprehensive study by Monthony et al. (2026) integrated over 130 RNA-seq libraries with ethylene pathway metabolite profiling and whole-genome sequencing across three XX and XY genotypes treated with STS and ethephon, respectively. This work demonstrated that ethylene-mediated sexual plasticity involves both systemic and local signaling components, where early transcriptional activation of ethylene biosynthesis and signaling genes occurred within 18 hours of treatment in leaves, prior to the emergence of flowers, while the new floral identity in developing flowers involved distinct sets of genes differentially regulated in each chromosomal sex. Other transcriptomic studies have similarly identified ethylene-related genes as candidates for sex determination (Adal et al., 2021), and network ontology analyses have further highlighted hormone-related gene expression changes associated with sexual plasticity (Orozco et al., 2025). Together, these findings have established ethylene as a primary hormonal driver of sexual plasticity in cannabis but also raised the question of whether other phytohormone pathways, particularly gibberellins, participate in this regulatory network.

The gibberellins (GAs) comprise a large family of tetracyclic diterpenoid phytohormones that regulate diverse aspects of plant growth and development, including stem elongation, flowering, fertility, and reproductive organ development (Bao et al., 2020; Pimenta Lange & Lange, 2006; Plackett et al., 2011; Wilson et al., 1992; Yamaguchi, 2008; Yu et al., 2004). Although more than one hundred gibberellins have been identified, only a small subset, primarily GA_1_ and GA_4_ in plants, are bioactive (Shani et al., 2024), with the remaining acting as precursors or inactive forms (Hedden & Thomas, 2012). *In planta* GA activity largely depends on the balance between GA production and deactivation (Yamaguchi, 2008). GA biosynthesis is a complex, multi-step process (Fig. 1) beginning with a 20-carbon precursor, geranylgeranyl diphosphate (GGPP), which undergoes enzymatic transformation to form GA_12_, the common precursor for all GAs in plants (He et al., 2020; Shani et al., 2024). From GA_12_, two parallel branches are commonly described: a non-13-hydroxylated route yielding GA_4_ and a 13-hydroxylated route in which GA_12_ is converted to GA_53_ and ultimately to GA_1_. In both branches, GA20-oxidases generate C19 precursors (e.g., GA_9_/GA_20_), and GA3-oxidases convert these into bioactive GAs (GA_4_/GA_1_) in the cytosol (Shani et al., 2024). GA deactivation is classically mediated by GA2-oxidases, which can also remove precursors from the biosynthetic pool.

**Figure 1.**
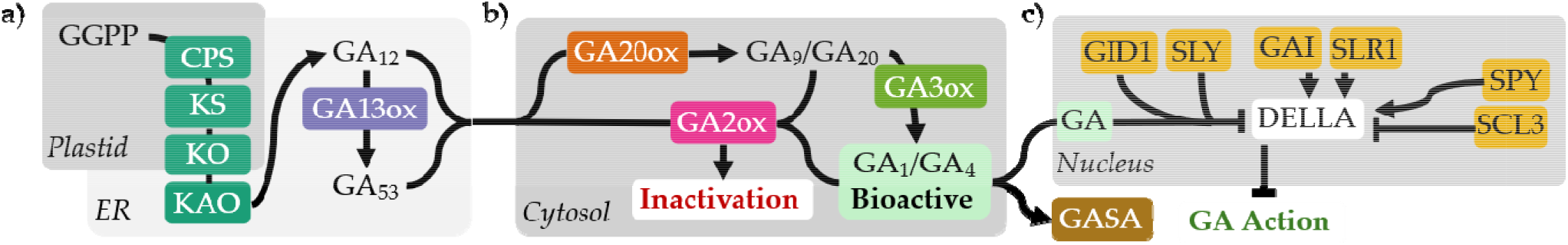
Canonical gibberellin (GA) biosynthesis, regulation and signaling pathway in plants. (a) GA early biosynthesis, converting geranylgeranyl diphosphate (GGPP) to GA_12_ via copalyl diphosphate synthase (CPS), ent-kaurene synthase (KS), ent-kaurene oxidase (KO), and ent-kaurenoic acid oxidase (KAO), occurring in the plastid and endoplasmic reticulum (ER), and 13-hydroxylation of GA_12_ by GA13-oxidase (GA13ox). (b) In the cytosol, GA_12/53_ are converted via parallel routes to the bioactive GAs GA_1_ and GA_4_ through the sequential activity of GA20-oxidases (GA20ox) and GA3-oxidases (GA3ox). Deactivation of bioactive GAs is catalyzed by GA2-oxidases (GA2ox). (c) GA signaling in the nucleus, where bioactive GA binds the receptor GID1, promoting formation of a GID1–GA–DELLA complex that recruits the F-box protein SLY1/SLY2 (SLY), leading to DELLA degradation repression of GA-responsive genes. DELLA activity is further modulated by SPY and SCL3. GAI and SLR1 represent DELLA-family transcriptional repressors. GA action induces GA-stimulated (GASA) genes.

Perception and signaling of bioactive GAs occur primarily in the nucleus, where GA binds to the soluble receptor GA INSENSITIVE DWARF1 (GID1) (Griffiths et al., 2007; Hirano et al., 2008) and promotes formation of a complex that triggers DELLA protein degradation via SLEEPY1 (SLY1) or its homolog SNEEZY (SNE/SLY2), both F-box subunits of SCF E3 ubiquitin ligase complexes (Ariizumi et al., 2011; McGinnis et al., 2003). DELLA proteins, belonging to the GRAS family, function as master growth repressors that integrate multiple phytohormone signals; their GA-dependent degradation represses downstream developmental programs (Davière & Achard, 2013). When GA activity is low, DELLAs accumulate and activate feedback mechanisms that modulate transcriptional regulation of GA metabolic genes, including GA20ox, GA3ox, and GA2ox, in several systems (Fukazawa et al., 2014; Hedden & Thomas, 2012). Downstream of these cores signaling events, GA-responsive genes are induced, including members of the GA-stimulated transcripts (GASA/GAST1-like; Snakin; Gibberellic acid-stimulated *Arabidopsis*) family, which encode small, secreted, cysteine-rich peptides that serve as integration nodes for multiple hormonal pathways (Aubert et al., 1998; Qu et al., 2016; Roxrud et al., 2007).

Beyond their roles in growth regulation, GAs serve as sex determinants across a range of plant lineages: in ferns and most eudicots, GAs generally promote male organ development, while in some monocots such as maize the effect is reversed (Gupta & Chakrabarty, 2013; Vandenbussche et al., 2007). In *Arabidopsis thaliana*, GA is essential for stamen filament elongation and anther development, and GA-deficient mutants display severe male sterility (Cheng et al., 2004; Plackett et al., 2011). In the dioecious relative spinach, the DELLA gene *SpGAI* (GIBBERELLIC ACID INSENSITIVE) shows gender-specific expression that is critical for unisexual organ initiation: high *SpGAI* levels in females repress B-class homeotic genes that promote maleness, while GA application induces *SpGAI* degradation, releasing this repression and causing masculinization (West & Golenberg, 2018). Other studies have also linked GA and sex determination in spinach (Ma et al., 2024; Wang et al., 2025; Y. Zhang et al., 2024).

In *C. sativa* specifically, a genome-wide association study identified a major sex determination QTL (*QTLSex_det1*) containing a gene encoding a GAI-like DELLA protein (Petit et al., 2020), suggesting that a similar DELLA-mediated mechanism may operate in cannabis (Salentijn et al., 2019). The link between gibberellins and sex expression in this species has been recognized since the pioneering work of Ram and Jaiswal (1972) and Chailakhyan (1979), who demonstrated that exogenous applications of GA_3_ could induce male flowers on genetically female plants. More recently, Alter (2024) showed that inflorescence development in female cannabis is mediated by photoperiod and GA: short-day conditions trigger a decrease in bioactive GA_4_ levels at the shoot apex, which is required for the formation of compact, cannabinoid-rich inflorescences, while exogenous GA_3_ application mimics long-day conditions and prevents condensed inflorescence formation. Transcriptomic analyses of sex-changed cannabis plants have also identified specific GA-related genes associated with sexual plasticity. In STS-induced male flowers on female plants, a putative gene encoding a GA 2-beta-dioxygenase (GA2ox) is consistently downregulated, suggesting reduced GA degradation and higher GA levels in masculinized tissues (Toscani et al., 2026). The gibberellin-regulated gene *GASA4* was found to be upregulated in both true male flowers and STS-induced male flowers compared to normal female flowers (Adal et al., 2021), and a cytochrome P450 gene suggested to be involved in GA biosynthesis was consistently upregulated in male tissues (Orozco et al., 2025).

Crucially, the GA and ethylene pathways do not operate in isolation. In model species, GA– ethylene crosstalk is mediated largely through DELLA proteins, which serve as integration hubs for both pathways. Ethylene signaling stabilizes DELLA proteins, thereby reducing GA-responsive growth and development (Achard et al., 2003, 2007). Activated ethylene signaling can also reduce levels of bioactive GAs by inhibiting GA biosynthesis enzymes such as GA20-oxidase (Achard et al., 2007), and the ethylene-regulated transcription factor EIN3 can physically interact with JAZ repressors, which themselves interact with DELLAs (Colebrook et al., 2014; Davière & Achard, 2013). In certain developmental contexts, ethylene and GA can act either antagonistically or synergistically: ethylene inhibits root elongation by blocking GA-induced DELLA degradation, while in floral induction GA promotes flowering and ethylene typically delays it (Achard et al., 2007; Sun, 2008). These molecular interactions provide a mechanism through which manipulation of one pathway could trigger downstream changes in the other, potentially explaining why ethylene-modulating treatments in cannabis lead to coordinated shifts in both ethylene- and GA-related gene expression.

Despite well-documented phenotypic effects of manipulating GA and ethylene pathways in C. sativa, the transcriptional basis of their crosstalk during sex reversal remains uncharacterized. Thus, GA-related genes (GARGs) make for strong candidates to understand the regulation associated with sex determination and sex expression. Building on the experimental framework established by the ethylene-focused analysis of Monthony et al. (2026), this study reconstructs the canonical GA pathway in *C. sativa* and examines expression patterns of key genes during these ethylene-driven sex reversals. By profiling CsGARGs expression across treated and untreated plants at multiple time points, we assess GA-related transcriptional shifts arising from ethylene– GA crosstalk, contributing to an integrative model of hormonal control of sex expression in cannabis. More broadly, elucidating this crosstalk has direct implications for the precise control of specialized metabolite production and the development of sex-stable genotypes, and positions *C. sativa* as a compelling model for investigating how GA-related pathways shape sexual plasticity in dioecious plants.

## Materials and Methods

### Identification of Orthologs of GARGs in *C. sativa*

A comprehensive literature review was undertaken to compile a list of GARGs in the canonical GA biosynthesis, regulation, and signaling pathways (Castro-Camba et al., 2022; Davière & Achard, 2013; Hedden, 2020; Hedden & Thomas, 2012; Sun & Gubler, 2004; Yamaguchi, 2008). Using the model species *Arabidopsis thaliana* L., a list of GARGs was used to identify *C. sativa* GARGs (CsGARGs). Inter-species orthologs were identified using OrthoFinder (Emms & Kelly, 2015) and BLASTP searches (Camacho et al., 2009) for unmatched genes of interest. For *C. sativa*, the reference genome ASM2916894v1 (cultivar ‘Pink Pepper’; RefSeq accession GCF_029168945.1) was used, alongside the fully phased haploid X and Y genomes from genotype 284 ‘Otto II’ (Carey et al., 2024) to improve orthogroup identification on sex chromosomes. The TAIR10.1 (RefSeq accession GCF_000001735.4) served as the *A. thaliana* reference. Nucleotide and protein sequences were retrieved from NCBI. The chromosome map was generated using chromoMap (v4.1.1) (Anand & Rodriguez Lopez, 2022) in R Studio (2025.5.1.513) running R (v4.5.0).

### Plant selection, flowering and sex plasticity induction

Vegetative plant propagation, growth conditions, chromosomal sex determination, flowering induction and sex plasticity treatments followed the procedures detailed by Monthony et al. (2026). Briefly, vegetative clones of a known sex from three *C. sativa* genotypes (La Rosca; *LR*, Panama Pupil V4; *PP*, Deadly Kernel; *DK*) were rooted, transferred to Pro-Mix BX substrate (Pro-Mix, Canada), and grown under controlled conditions (Conviron, Canada) with an initial 12/12 photoperiod, then shifted to 18/12 for flowering. Light intensity from LED lightning reached up to 750 µmol/m^2^/s, and substrate pH was maintained between 5.5 and 6.0.

Sex reversal in XX plants was induced using a 3 mM silver thiosulfate (STS) solution with 0.1% Tween 20, applied as foliar spray (~50 µL/plant) weekly for three weeks starting at the photoperiod shift (day 0). XY plants received a single application of 500 mg/L ethephon (diluted from 40% stock) with 0.1% Tween 20 at day 0. In both cases, untreated control plants were sprayed with distilled water containing 0.1% Tween 20 to ensure comparable conditions.

### RNA sequencing and transcriptome analysis

RNA extraction, library preparation, sequencing, and read processing steps are described in details in Monthony et al. (2026). Mature leaf tissues were harvested at day 0 and 1 (18 hours after treatment), and immature flowers were harvested at day 14. These samples were flash-frozen, ground, and total RNA was extracted using the RNeasy Plant Mini Kit (Qiagen GmbH, Germany). RNA quality was assessed by NanoDrop and Bioanalyzer, and samples with high integrity were used for cDNA synthesis with the NEBNext® Ultra™ II Directional RNA Library Prep Kit (New England Biolabs, Ipswich, MA, USA). Sequencing was performed at the *Institut de Biologie Intégrative et des Systèmes* (IBIS; Université Laval) on the Element AVITI platform, producing 150 bp paired-end reads (~30 million reads/sample). Reads were quality-checked with FastQC and trimmed using Trimmomatic. Contaminant sequences (e.g., animal, fungal RNA) were removed by BLAST filtering using a custom Trimmomatic pipeline. Cleaned reads were aligned to the *C. sativa* ‘Pink Pepper’ genome using STAR (v2.7.11b), gene-level quantification was performed with HTSeq-count (v2.0.2) and mapping quality was verified with Qualimap (v.2.2.1; Okonechnikov et al., 2015).

### Differential expression analysis and gene expression visualisations

Differentially expressed CsGARGs were identified using DESeq2 (v1.42.1; Love et al., 2014). Count data were filtered with HTSFilter and normalized using variance stabilizing transformation (VST). For day 0 and day 1 samples, correction was applied as described in Roy et al. (2026). Log_2_fold changes (LFC) were shrunk using the *ashr* method to improve estimates for low-expression genes. Genes with adjusted p-value ≤ 0.05 were considered significantly differentially expressed. Results were visualized using bar plots of LFC and boxplots of normalized counts for group comparisons using the R packages ggplot2 (v3.4.1; Wickham, 2009). The VST normalized, batch-corrected counts of genes of interests were represented through principal components analysis (PCA) plots using ggplot2 (v3.5.1) (Wickham, 2016). All analyses were performed in RStudio (2025.5.1.513) running R (v4.5.0).

## Results

### Primary GARGs are present in the C. sativa genome

Orthology-based searches identified 50 putative CsGARGs, of which 46 were expressed above the filtering threshold in the transcriptomic dataset and are summarized in Table 1. The gene collection is publicly available on the National Center for Biotechnology Information website (NCBI) (Bethesda, MD) (https://www.ncbi.nlm.nih.gov/sites/myncbi/julien.roy.2/collections/63974920/public/).

This gene set covers the major enzymatic, regulatory, and signaling components of the canonical GA pathway. Putative orthologs were identified for the early biosynthetic steps leading to GA_12_, including CPS, KS, KO and KAO genes, corresponding to CsGA1, CsGA2, CsGA3, and two CsKAO paralogs (CsKAO1, CsKAO2) (Fig. 2a). Downstream of GA_12_, multiple paralogs were identified in the three oxidase families controlling GA activation and inactivation, namely GA20ox, GA3ox, and GA2ox. The GA3ox group included CsGA3ox1 and the CsGA3ox3 series, with four closely related paralogs, CsGA3ox3A–D, including three neighboring genes. Additional GA-modifying enzymes were also identified, including one putative CYP714A gene (CsCYP714A1) and four putative GA13ox/CYP72A9-like genes (CsCYP72A219A–D), supporting a reconstructed pathway that includes a GA13ox step connecting GA_12_ to GA_53_.

**Figure 2.**
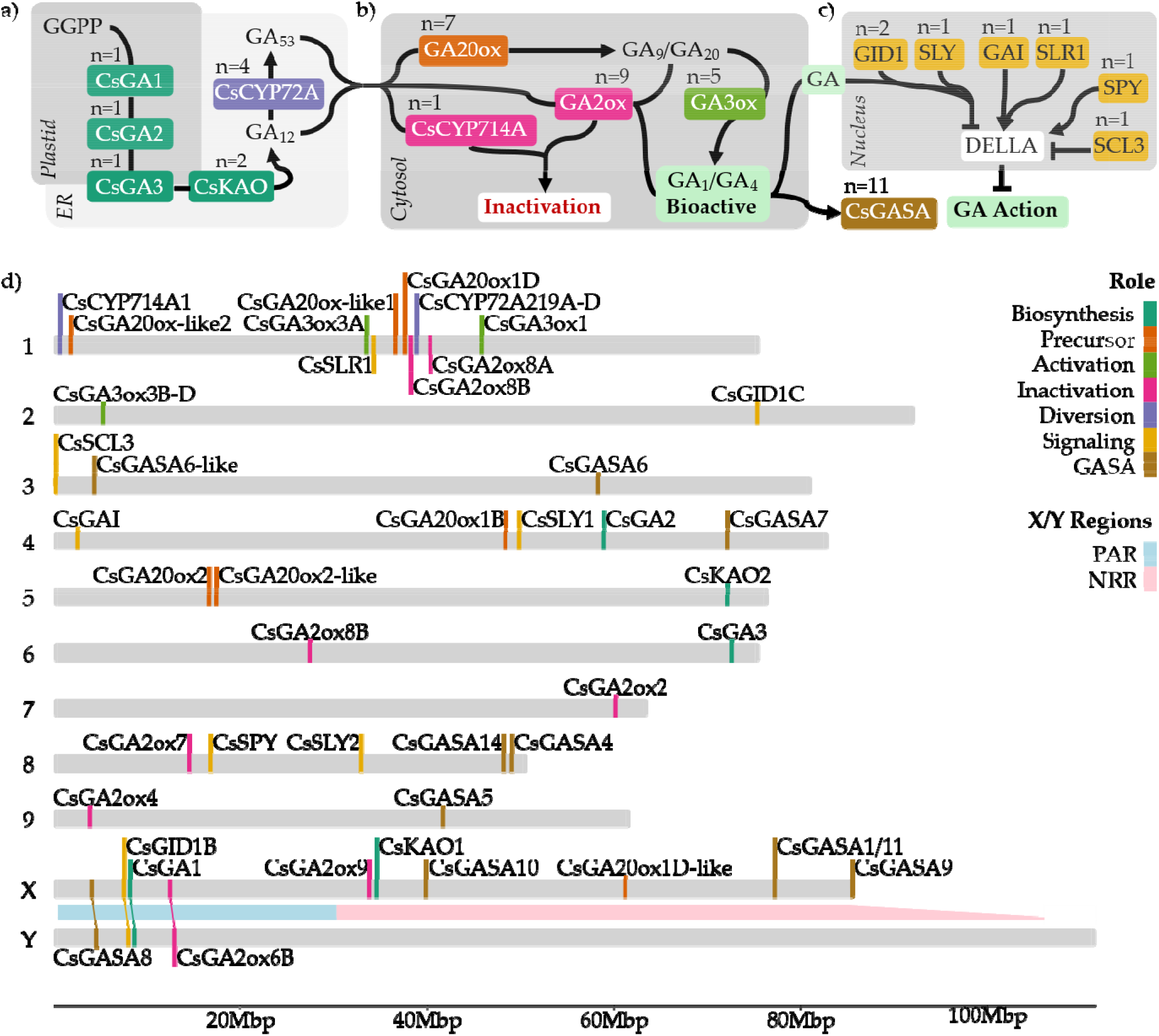
GA pathway and genomic localization of putative GARGs in C. sativa. a) Proposed GA biosynthesis and signaling pathway in C. sativa based on orthology with A. thaliana, showing enzymes involved in each step and the number of orthologs found. b) Chromosomal positions of CsGARGs across autosomes and sex chromosomes X/Y, color-coded by functional role. Autosomal gene positions are based on the ‘Pink Pepper’ reference genome; Pseudo-autosomal region (PAR) and non-recombining regions (NRR) on the X and Y chromosomes follow Carey, Bentz, et al. (2024).

Core GA signaling components were also mapped, including two putative GA receptors (CsGID1B, CsGID1C), DELLA-related genes (CsGAI, CsSLR1) and the signaling regulator CsSCL3, as well as the F-box genes CsSLY1 and CsSLY2/SNE, which are involved in DELLA degradation. The dataset also included the negative regulator CsSPY and multiple GASA genes representing GA-responsive outputs.

Genomic localization of these CsGARGs revealed a widespread distribution across the *C. sativa* genome (Fig. 2d). The X chromosome harbored the highest number of CsGARGs (11), five of which were in the pseudoautosomal region of the sex chromosomes. No genes were detected in the non-recombinant region (NRR) of the Y chromosome, whereas six were located within the X chromosome NRR.

### Transcriptome results

Following filtration of low-count genes, 34 of the 50 CsGARGs remained in leaf samples (day 0 and day 1). Principal component analysis (PCA) of CsGARGs expression revealed time-dependent structuring of samples according to floral phenotype (Fig. 3a-c). Baseline (day 0) analysis of GARGs shows broad similarity in expression profiles during vegetative growth (Fig. 3a), with PC1 and PC2 explaining 47% and 14% of the variance, respectively. Immediately following treatment and photoperiod change (day 1), expression profiles show perturbation, but do not cluster by chromosomal sex (XX vs XY) or by sexual phenotype class, as indicated by the overlap of the 95% ellipses (Fig. 3b), with PC1 and PC2 explaining 58% and 10% of the variance, respectively.

**Figure 3.**
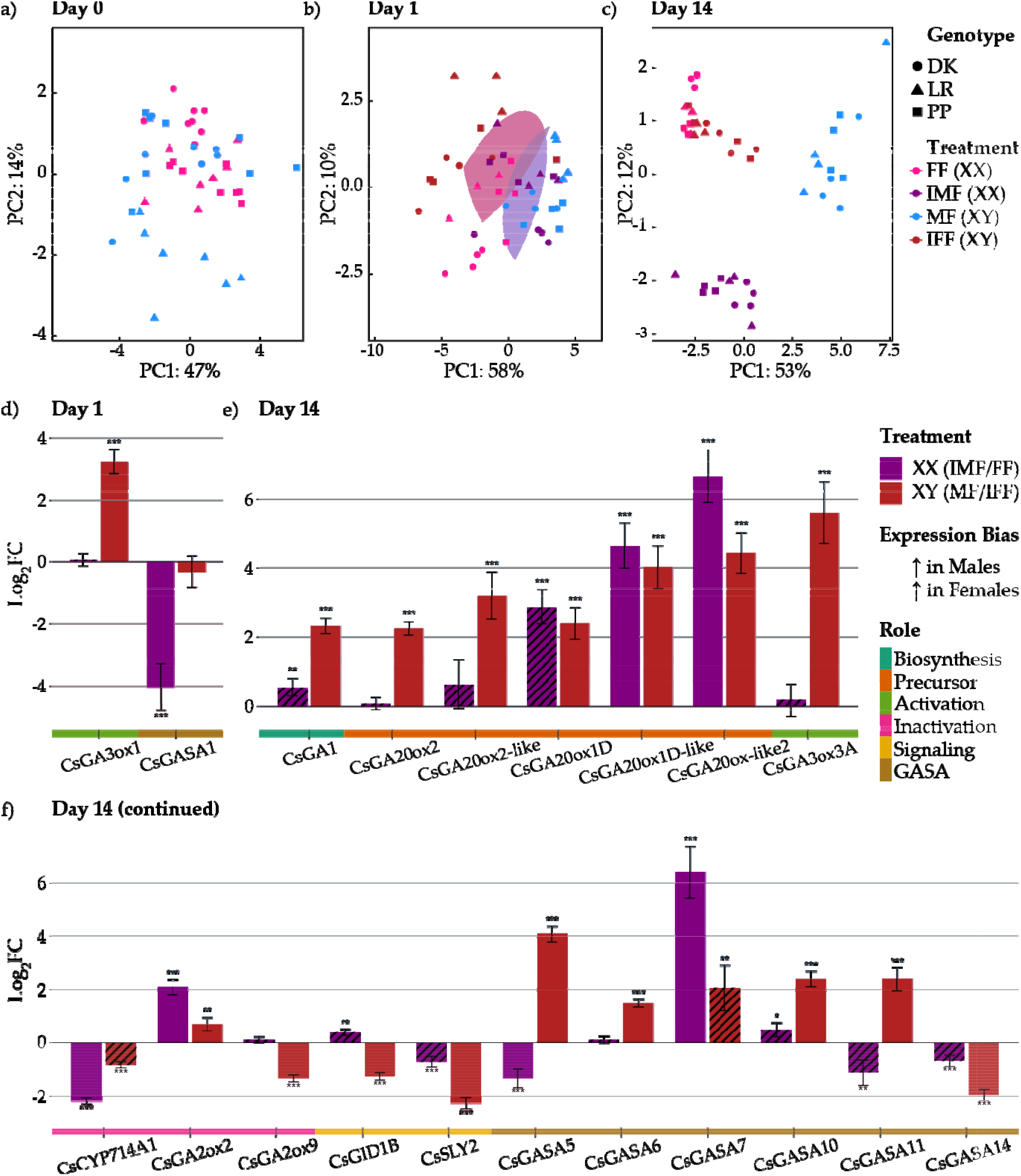
CsGARG expression profiles across treatments and timepoints. (a–c) Principal component analysis (PCA) of batch-corrected variance stabilized transformed (VST) counts of CsGARGs. (a) Day 0 leaf samples (n = 34 genes); (b) Day 1 leaf samples (n = 34 genes); (c) Day 14 immature flower samples (n = 45 genes). Each point represents an individual sample; ellipses show 95% confidence intervals for each treatment group. Colors indicate treatment groups and symbols represent genotypes (DK = Deadly Kernel; LR = La Rosca; PP = Panama Pupil V4). (d) Differential expression of CsGARGs at day 1 following ethylene pathway manipulation, grouped by functional role. Bars represent log_2_ fold changes (LFC) relative to untreated controls for silver thiosulfate (STS) treatment on genotypically female plants (XX; IMF vs. FF, purple) and ethephon treatment on genotypically male plants (XY; MF vs. IFF, red). (e–f) Differential expression of CsGARGs at day 14, grouped by functional role, for the STS contrast (XX) and ethephon contrast (XY), respectively. Only genes with adjusted *p*-value ≤ 0.05 in all genotypes and |LFC| ≥ 1 are displayed. Asterisks denote significance levels (\*\**padj < 0.05; **padj < 0.01; ***padj* < 0.001).

At day 14, 45 of the 50 CsGARGS were expressed in immature flower samples following low-count filtration. At this stage, the PCA shows a more structured pattern of clustering (Fig. 3c). FF (XX) and IFF (XY) samples overlap despite their different sex chromosome karyotypes, indicating clustering by sexual phenotype rather than genotype, with PC1 and PC2 explaining 53% and 12% of the variance, respectively. In contrast, phenotypic male samples form distinct groups: IMF (XX) does not overlap with MF (XY). Moreover, IMF (XX) samples are clearly separated from their untreated counterparts (FF), indicating a shift associated with ethylene inhibition and the acquisition of a male phenotype.

Only two CsGARGs responded in leaves within 18 hours of sex plasticity induction (Day 1). In XY plants, CsGA3ox1 was significantly downregulated (log_2_FC = −3.24, adjusted *P*-value = 1.26e-15) following ethephon treatment compared to control (Fig. 3c). In XX plants treated with STS, CsGASA1 was significantly downregulated (log_2_FC = −4.04, p = 2.18e-7) relative to controls XX (Fig. 3d).

At day 14, several GARGs were differentially expressed across treatment groups (Fig. 3e). In the ethephon contrast (MF vs IFF; XY), *CsGA1* showed significantly higher expression in untreated immature flowers (log_2_FC = 2.33, adjusted *P*-value = 1.98e-23). Multiple GA20-oxidase paralogs (*CsGA20ox2, CsGA20ox2-like, CsGA20ox1D, CsGA20ox1D-like* and *CsGA20ox-like2*) were also significantly more expressed in untreated samples indicating higher expression in phenotypic male flowers, with log2FC between 2.25 and 4.43. A GA3-oxidase gene shows strong downregulation following treatment, with higher expression in control males (log_2_FC = 5.61, adjusted *P*-value = 3.19e-11). In contrast, expression of CsSLY2 was significantly lower in untreated males (log_2_FC = −2.27, adjusted P-value = 8.17e-25). Members of the GASA gene family display strong responses, with many paralogs strongly downregulated by treatment (CsGASA5, CsGASA6, CsGASA10, CsGASA11), while CsGASA14 showed lower counts in control plants (log_2_FC = −1.96, adjusted *P*-value = 3.47e-21).

In the STS contrast (IMF vs FF; XX), four genes were significantly differentially expressed at day 14, including CsGA20ox1D-like (log_2_FC = 4.64, adjusted *P*-value = 2.25e-12) and CsGA20ox-like2 (log_2_FC = 6.66, adjusted *P*-value = 6.00e-18), which showed the same phenotype-associated pattern observed in the ethephon contrast, with higher expression in phenotypic male flowers. CsCYP714A1 was significantly downregulated by treatment (log2FC = −2.19, adjusted P-value = 9.84e-63) in comparison to untreated XX. Finally, CsGASA7 was significantly upregulated in IMF (log2FC = 6.40, adjusted P-value = 4.75e-11).

## Discussion

### Early dynamics of the GA response during sex reversal

A feature of phytohormone signaling is a plant’s ability to exert temporal control over the pathway to adjust biological outcomes in responses to internal and external stimuli. In the present study we observed GA-related transcriptional responses to ethylene-induced sex change within 18 hours of treatments. *CsGA3ox1*, the sole GA3ox expressed in leaves at day 1, was strongly downregulated in ethephon-treated XY plants (IFF), suggesting that ethephon treatment rapidly attenuates bioactive GA production, potentially facilitating the hormonal shift required for feminization. Notably, *CsGA3ox1* was not differentially expressed at day 1 in the STS treatment, an asymmetry that may reflect the differing nature of the two treatments: ethephon directly increases ethylene levels through a single application, whereas STS acts as an inhibitor of ethylene perception and requires repeated applications over three weeks to achieve sex reversal. The cumulative nature of the STS treatment may result in a gradual perturbation of the GA pathway, with transcriptional changes in GA biosynthetic genes emerging later or below the detection threshold after the first application.

In contrast, STS treatment elicited downregulation of *CsGASA1*, a GA-responsive output gene, suggesting the two treatments engage the GA pathway at different levels, consistent with their distinct modes of action. These results suggests that the broader GA changes observed in developing flower tissues may be initiated via *CsGA3ox1* and *CsGASA1* in vegetative tissues via ethylene signaling. This limited early response contrasts with the broader ethylene-related transcriptional activation reported at the same time point by Monthony et al. (2026), where multiple ethylene biosynthesis and signaling genes responded within 18 hours of treatment. Nevertheless, the rapid recruitment of GA-responsive genes within 18 hours of treatment, in vegetative leaf tissue and prior to any visible floral transition, implies that ethylene-GA crosstalk is engaged quickly upon hormonal perturbation. This early coordination suggests that the two pathways may be coupled at the onset of the sexual transition, pointing to an signaling network initiating sexual plasticity before morphological changes are apparent.

### CsGA1 and multiple GA20ox genes control GA activity in developing flowers

The downregulation of *CsGA1* in ethephon-treated XY plants is consistent with the broader suppression of GA biosynthetic capacity observed in feminized plants. *CsGA1* is responsible for the first committed step in GA biosynthesis (Hedden, 2020). Its higher expression in untreated male flowers suggests that sustained GA biosynthetic flux contributes to the maintenance of the male floral development, and that ethephon-driven feminization involves attenuation of this flux at its earliest committed step. Together with the downregulation of *CsGA3ox1* at day 1, the reduction of *CsGA1* at day 14 suggests that ethylene may suppress GA production broadly by reducing the availability of early precursors, leading to a progressive reduction of GA activity accompanying the transition to a female floral phenotype.

Coordinated changes across multiple GA20-oxidase orthologs were found in developing flowers, all of which showed higher expression in phenotypic male flowers relative to phenotypic females. As GA20ox enzymes convert substrates into precursors to bioactive GAs (He et al., 2020; Hedden & Thomas, 2012), their preferential expression in phenotypic male flowers regardless of chromosomal sex is consistent with elevated GA flux driving masculinization. On the other hand, the downregulation of *GA20ox* orthologs in IFF relative to untreated males is consistent with reduced GA biosynthetic capacity during feminization.

Notably, the genomic localization of *CsGA20ox1D-like* on chromosome X (position ~61.4 Mb), places it within a region that has been proposed to be subject to sex-chromosome-specific regulatory dynamics (Toscani et al., 2025). The presence of a GA biosynthetic gene on the X chromosome raises the possibility that GA pathway components may be subject to sex-linked regulatory influences.

The downregulation of *CsGA3ox3A* in IFF (XY) immature flowers, with higher expression maintained in untreated phenotypic males, is consistent with the broader suppression of GA biosynthetic capacity accompanying ethephon-driven feminization. Notably, this ortholog shows comparatively low overall expression in our dataset relative to the predominantly expressed *CsGA3ox1*, and its expression is effectively restricted to XY plants under control conditions, with ethephon-treated XY plants showing expression reduced to near zero. Although *CsGA3ox1*, significantly downregulated in leaf tissue of IFF plants, did not meet the log2FC threshold, it should be noted that the expression level in immature flowers was significantly higher in IMF (XX) (log_2_FC = −0.49, adjusted *P*-value = 0.006) when compared to control females (FF).

The upregulation of two *CsGA2ox* orthologs in opposing sex-reversal contexts, with one induced in IMF (XX) plants and another in IFF (XY) plants, points to differential responses to treatments. GA2ox enzymes deactivate bioactive GAs and their immediate precursors, serving as the principal mechanism by which plants reduce active GA pools (Hedden & Thomas, 2012; Yamaguchi, 2008). The engagement of distinct paralogs in each treatment context suggests that GA homeostasis during sex reversal is not achieved through a single deactivation route, but rather through paralog-specific programs tuned to the shift in floral identity. This pattern aligns with broader observations that *GA2ox* family members are often functionally specialized, with individual paralogs exhibiting distinct tissue specificities, substrate preferences, and developmental roles (Rieu et al., 2008; Shani et al., 2024).

The location of *CsGA2ox9*, upregulated in IFF (XY) maps to the NRR of the X chromosome. Its preferential expression in feminized XY plants, which carry only a single X chromosome, could indicate that this paralog has acquired sex-biased regulatory features through its location in the NRR. Functionally, induction of *CsGA2ox9* in feminized XY plants would be expected to accelerate deactivation of bioactive GAs, reinforcing the suppression of the masculinizing GA biosynthetic program already evidenced by downregulation of *CsGA1* and *CsGA20ox* paralogs. Conversely, induction of a *CsGA2ox2* in masculinized XX plants may reflect compensatory feedback regulation or pleiotropic engagement of GA2ox in processes unrelated to bioactive GA pool size. Resolving these possibilities will require paralog-specific functional characterization and direct measurement of bioactive GA levels in sex-reversed tissues.

The downregulation of *CsGA1*, multiple *CsGA20ox* orthologs, and the GA3ox/GA2ox dynamics in IFF samples indicate that ethephon-driven feminization engages a multi-tiered suppression of GA activity spanning biosynthesis, precursor formation and final activation. Such transcriptional repression of GA biosynthesis is consistent with the *Arabidopsis* model in which ethylene signaling reduces bioactive GA content through attenuation of multiple biosynthetic steps (Achard et al., 2007), and it suggests that ethylene-driven sex reversal in *C. sativa* operates, at least in part, by dismantling the GA biosynthetic program that supports the male floral identity.

Beyond sex determination, elevated GA biosynthetic activity in phenotypic male flowers may also underlie the architectural differences between male and female *C. sativa* plants. Male plants undergo pronounced internode elongation during the floral transition, producing taller, more racemose inflorescences with greater internodal spacing compared to the compact, condensed inflorescences characteristic of female plants. Alter et al. (2024) demonstrated that GA4 levels in the shoot apex are inversely correlated with inflorescence condensation in female *C. sativa*, with elevated GA promoting internode elongation under short-day conditions. The preferential expression of GA20ox paralogs and *CsGA1* in phenotypic male flowers observed here may contribute not only to male sex organ development but also to the elongated inflorescence of staminate plants, suggesting a role in sexual identity and broader morphological dimorphism between sexes.

### GA signaling adjustments: *CsCYP714A1, CsGID1B* and *CsSLY2*

Beyond biosynthesis, our results revealed significant changes in GA signaling. *CsCYP714A1* was significantly downregulated in IMF relative to untreated female controls at day 14. In Arabidopsis, *CYP714A1* encodes a cytochrome P450 monooxygenase that divert GA_12_, the key precursor of all GAs, away from the biosynthetic pool and toward inactive forms: its overexpression produces severe GA-deficient dwarfism (Nomura et al., 2013; Y. Zhang et al., 2011). The downregulation in IMF is therefore consistent with a metabolic shift that preserves GA_12_ availability, allowing greater flux into the biosynthetic pathway to support the elevated *GA20ox* expression and, presumably, higher bioactive GA levels associated with the male phenotype.

Conversely, *CsGID1B* is upregulated in IFF (XY). GID1 proteins are the soluble GA receptors responsible for perceiving bioactive GA and initiating DELLA degradation via the SCF-SLY1 ubiquitin-proteasome pathway (Griffiths et al., 2007; Hirano et al., 2008). Upregulation of a GA receptor in a context where GA biosynthesis is suppressed appears paradoxical. However, it may reflect a feedback mechanism: when bioactive GA levels fall and DELLA proteins accumulate, the GA signaling response is upregulated as part of a homeostatic reaction to restore GA perception capacity (Hedden & Thomas, 2012). Under this interpretation, *CsGID1B* induction in IFF plants would be a consequence of the reduced GA biosynthetic output. Alternatively, *CsGID1B* upregulation may serve a more functionally specific role during female floral development. GID1 receptors are active participants in shaping the sensitivity GA signaling, with different GID1 paralogs exhibiting distinct affinities for bioactive GA forms and differential interactions with specific DELLA proteins (Nakajima et al., 2006). Induction of CsGID1B in feminized flowers could therefore reflect a reconfiguration of GA signaling sensitivity rather than simply a homeostatic compensation, potentially tuning the tissue to respond differently to residual or locally imported GA. This interpretation is consistent with the broader GA biology framework in which spatial and temporal control of receptor expression, rather than GA biosynthesis alone, shapes the developmental output of GA signaling (Binenbaum et al., 2018).

A third possibility is that *CsGID1B* upregulation is linked to ethylene signaling. In *Arabidopsis*, ethylene modulates DELLA stability through CTR1-EIN3-dependent pathways, and the resulting changes in DELLA accumulation feed back to regulate both GA biosynthetic and signaling genes (Achard et al., 2003, 2007). If ethephon treatment in XY plants drives DELLA accumulation by reducing bioactive GA, the induction of *CsGID1B* may reflect this feedback rather than a direct transcriptional response to ethylene.

*CsSLY2* (SNEEZY), encoding an F-box protein that targets DELLA repressors for proteasomal degradation, thereby relieving repression of GA-responsive programs, was upregulated in induced female flowers at day 14 (Ariizumi et al., 2011; McGinnis et al., 2003). In *A. thaliana, SLY2* targets a subset of DELLA proteins (*RGA* and *GAI*) but does not regulate *RGL2*, distinguishing it functionally from its homolog *SLY1*. The upregulation of *CsSLY2* during feminization of XY individuals is intriguing because it would be expected to enhance DELLA degradation, potentially activating specific GA-responsive programs even as overall GA biosynthetic capacity (reflected by *GA20ox* expression) decreases. Specific developmental outputs dependent on SLY2-mediated DELLA clearance may be selectively activated while bulk GA production is attenuated.

### GASA gene family members as downstream integrators of hormonal crosstalk

The GASA (Gibberellic Acid Stimulated *Arabidopsis*) gene family emerged as one of the most transcriptionally dynamic components of the GA response during sex reversal. Multiple GASA paralogs (*CsGASA5, CsGASA6, CsGASA10, CsGASA11*) were strongly downregulated in IFF relative to controls, while *CsGASA14* was significantly upregulated. In IMF, *CsGASA7* showed strong upregulation. GASA proteins are small, secreted, cysteine-rich peptides that function as downstream effectors of GA signaling and as integration nodes for multiple hormonal pathways (Aubert et al., 1998; Roxrud et al., 2007). In *Arabidopsis*, GASA family members play diverse roles in reproductive development: *GASA4* positively regulates floral meristem identity and the transition to flowering, *GASA5* acts as a negative regulator of GA-induced flowering and stem growth, and *GASA14* promotes growth and leaf expansion while modulating reactive oxygen species accumulation (S. Zhang et al., 2009). Critically, GASA genes are known to integrate signals from multiple hormonal pathways, as several family members, including *GASA4* and *GASA6*, are induced by GA, brassinosteroids, and auxin but repressed by stress-associated hormones such as abscisic acid, jasmonic acid, and salicylic acid (Qu et al., 2016). This multi-hormonal responsiveness positions GASA genes as potential nodes of ethylene-GA crosstalk, guiding developmental outcomes. The observation that GASA expression patterns track with floral phenotype rather than chromosomal sex during sex reversal suggests that these peptides may serve as functional readouts of the hormonal state of the developing flower. Specifically, the broad downregulation of multiple GASA paralogs in IFF suggests a coordinated suppression of GA-responsive outputs during feminization, consistent with the reduced GA20ox expression observed in the same samples and with the established role of GA as a promoter of male flower development in *C. sativa*.

Notably, several GASA genes are located on the X chromosome (Fig. 2d) : *CsGASA1, CsGASA8, CsGASA9, CsGASA10*, and *CsGASA11*. Of these, *CsGASA8* resides in the PAR, while the remaining four are located within the NRR of the X chromosome, with *CsGASA1* and *CsGASA11* in close physical proximity (~77.2 Mb), immediately upstream of the monoecy locus recently identified by Carey et al. (2024), raising the possibility that this chromosomal region concentrates regulatory elements at the intersection of GA and ethylene signaling. The early downregulation of *CsGASA1* following STS treatment in XX plants may therefore reflect rapid ethylene perturbation at a sex-chromosome-linked locus, analogous to the behavior of X-linked ERGs such as *CsACO5* and *CsEBF1* reported by Monthony et al. (2026), and raises the possibility that this chromosomal region concentrates regulatory elements at the intersection of GA and ethylene signaling during sex determination.

### Evidence for GA–ethylene crosstalk during sexual plasticity

These findings extend a body of work in model species that has progressively positioned the DELLA proteins as central integrators of ethylene and GA signaling. In *Arabidopsis*, Achard et al. (2003) demonstrated that ethylene inhibits root growth and modulates apical hook development by stabilizing DELLA repressors through a *CTR1*-dependent pathway, with the ethylene-induced delay in GA-mediated DELLA degradation providing a direct molecular link between the two hormones. Subsequent work showed that this DELLA-centered crosstalk extends to the floral transition: activation of ethylene signaling reduces bioactive GA levels, enhances DELLA accumulation, and consequently represses the floral meristem identity genes *LFY* and *SOC1*, delaying flowering particularly under non-inductive short-day photoperiods (Achard et al., 2007). The mechanism operates through *CTR1*-*EIN3*-dependent signaling, with *ERF1* acting downstream to promote DELLA accumulation by reducing GA content. Achard et al. (2004) further uncovered a downstream layer of regulation in which GA-DELLA signaling modulates miR159 levels, which post-transcriptionally regulates GAMYB transcription factors implicated in both floral initiation and anther development, linking hormonal status to floral developmental programs.

The integration of ethylene and GA signaling appears to be broadly conserved across developmental contexts, with DELLA proteins serving as a hub for hormonal cross-regulation in vegetative growth, apical hook formation, hypocotyl elongation, and root development (Van De Poel et al., 2015). The recurring theme across these systems is that ethylene modulates GA biosynthesis, GA-mediated DELLA degradation, or downstream GA-responsive outputs, often in a tissue- and context-specific manner. Our observations in *C. sativa* are consistent with this framework: ethephon treatment downregulated *CsGA3ox1* at day 1 and *CsGA1* at day 14, suggesting progressive attenuation of bioactive GA production, while broad downregulation of *GASA* paralogs in feminized plants points to coordinated suppression of GA-responsive outputs. These shifts align with the Arabidopsis model in which elevated ethylene signaling reduces GA biosynthetic capacity and dampens downstream GA-responsive transcription (Achard et al., 2007), and extend the developmental consequence from delayed flowering to reversal of floral sexual identity.

The ethylene-GA crosstalk operates through multiple regulatory layers beyond the DELLA hub. In Arabidopsis, ethylene treatment broadly downregulates members of the *GA20ox* and *GA3ox* families, while exogenous GA upregulates several *ACS* genes, indicating bidirectional control at the level of gene expression (Dugardeyn et al., 2007). Analysis of the *gai eto2-1* double mutant revealed reciprocal post-translational control between the two pathways: GA signaling modulates ACS5 protein stability through an ETO1-independent degradation pathway, while ethylene may in turn influence DELLA turnover by affecting the DELLA-SLY1/2 interaction (De Grauwe, Chaerle, et al., 2008; De Grauwe, Dugardeyn, et al., 2008). The DELLA proteins themselves control GA homeostasis by activating transcription of GA biosynthetic genes (*GA20ox, GA3ox*) through the DELLA-GAF1 complex under GA-deficient conditions (Fukazawa et al., 2014). In the presence of high GA levels, DELLA is degraded via GID1-SLY1/SLY2, creating a negative feedback loop (Hedden & Thomas, 2012). Our observation that ethephon treatment (which promotes ethylene signaling) is associated with reduced *GA20ox* expression and a reorganized *GASA* profile in IFF is consistent with the DELLA-stabilizing effect of ethylene, which is expected to suppress GA biosynthetic gene expression, while potentially activating specific signaling outputs through SLY2-mediated selective DELLA degradation.

The link between ethylene-GA crosstalk and sex determination has precedent beyond cannabis. In dioecious *Spinacia oleracea*, GA application masculinizes XX plants while paclobutrazol feminizes XY plants, and the DELLA protein *SpGAI* was identified as the feminizing factor acting upstream of B-class floral identity genes (West & Golenberg, 2018). Notably, ethylene also influences spinach sex expression, and the convergence of GA-DELLA and ethylene signaling on B-class gene regulation parallels the integrated regulatory architecture we observe in *C. sativa*. Similarly, in *Cucumis melo* and other cucurbits, ethylene biosynthesis genes (ACS) act as primary sex determinants, and GA modulation can shift sexual phenotype, suggesting that ethylene-GA crosstalk may represent a recurrent regulatory motif in the hormonal control of plant sex expression (Boualem et al., 2015).

*C. sativa* is therefore a valuable system for dissecting how ethylene-GA crosstalk is wired into sex determination in dioecious plants, and they suggest that the broader principles emerging from other angiosperms, namely that DELLA proteins integrate ethylene and GA signals to govern reproductive fate, are likely operative in cannabis as well. In ethylene inhibited systems, such as during STS treatment, the resulting masculinization of XX plants was associated with enhanced GA20ox expression and downregulation of *CsCYP714A1*, conditions which are required to support increase in GA biosynthetic flux, such as has been demonstrated in Arabidopsis (Nomura et al., 2013; Rieu et al., 2008). This is consistent with the classical observation that GA application promotes male flower development in cannabis (Galoch, 1978; Ram & Jaiswal, 1972) and aligns with data from other dioecious species.

### Toward an integrative model of hormonal control of sexual plasticity

*C. sativa* sex reversal has largely been studied using ethylene-centered molecular models (Garcia-de Heer et al., 2025, 2026; Lubell & Brand, 2018; Monthony et al., 2026; Moon et al., 2020b; Orozco et al., 2025), largely overlooking the broader interconnectedness of plant growth regulatory pathways. The present study expands this framework by integrating gibberellins, a hormone family with a demonstrated role in *C. sativa* sexual plasticity, into hormonal events underlying sex reversal. Our results suggest that GA-mediated transcriptional restructuring is a downstream consequence of ethylene signaling, with GA-related responses remaining largely quiescent in the first 18 hours following chemical perturbation of endogenous ethylene biosynthesis, with only *CsGA3ox1* and *CsGASA1* responding. As the developmental program unfolds over the following two weeks, the GA pathway undergoes broad transcriptional restructuring that tracks with the acquisition of opposite-sex floral identity. This restructuring encompasses biosynthetic capacity (*CsGA1*, multiple *GA20ox* paralogs, *CsGA3ox3A*), precursor channeling (*CsCYP714A1*), inactivation (*CsGA2ox2* and *CsGA2ox*), signaling (*CsGID1B* and *CsSLY2*), and downstream output (the *GASA* gene family), suggesting comprehensive reorganization of the GA network as an integral component of the sex reversal process.

This temporal hierarchy is consistent with known mechanistic links between these pathways in model species. Stabilization of DELLA proteins by ethylene would initially suppress GA-responsive programs while simultaneously upregulating GA biosynthetic gene expression through homeostatic feedback (Achard et al., 2003; Fukazawa et al., 2014). As DELLA-mediated suppression relaxes during the transition to the new sexual phenotype, the resulting shift in GA regulation would then drive the morphological realization of the opposite-sex floral program. Whether GA signaling in turn feeds back onto the ethylene pathway remains to be explored in cannabis, though such bidirectional crosstalk is well documented in *Arabidopsis* (De Grauwe et al., 2007).

The interpretation of CsGARG transcriptional dynamics warrants consideration of the spatial complexity inherent to GA biology. GA is not a stationary signal as both bioactive forms and their precursors move within and between tissues, and the site of biosynthesis frequently differs from the site of action (Binenbaum et al., 2018). Expression measured in tissue therefore reflects local transcriptional states that do not necessarily indicate where bioactive GA is ultimately produced or perceived. Furthermore, the well-documented feedback regulation of GA biosynthesis genes by DELLA accumulation means that elevated *GA3ox* or *GA20ox* transcript levels can reflect low bioactive GA rather than high biosynthetic output (Hedden & Thomas, 2012).

The inherent pleiotropy of GA signaling also warrants caution in attributing shifts in expression exclusively to sex reversal. GA regulates stem elongation, inflorescence architecture, trichome development, and flowering in *C. sativa* (Alter et al., 2024), and some transcriptional changes observed here likely reflect the broader developmental reorganization. Disentangling sex-specific from sex-independent GA responses will require direct quantification of endogenous GA levels during sex reversal and functional perturbation of specific genes. The framework established here is well-positioned to inform these approaches.

## Conclusion

This study provides the first systematic characterization of the GA pathway in *C. sativa*, and direct transcriptomic evidence that ethylene perturbation reshapes GA-related gene expression during sex reversal in *C. sativa*, advancing an understanding of ethylene-GA crosstalk in dioecious plants. By demonstrating that ethylene manipulation triggers transcriptional reprogramming of CsGARGs, our results reveal a GA response that progresses from gene-specific early shifts to broad transcriptional remodeling during flower development. This sex-reversal leads to changes in GA activity, through *CsGA1* and multiple *GA20ox* genes, GA signaling, through Cs*CYP714A1, CsGID1B* and *CsSLY2*, and downstream regulation, through multiple *GASA* genes which track the acquisition of opposite-sex floral phenotypes. Ethylene-driven sex reversal in *C. sativa* engages GA signaling as an integral downstream module and the convergence of GA- and ethylene-related expression profiles broaden the model of sexual plasticity into the GA signaling domain. The enrichment of GA-related genes on the X chromosome further connects the GA signaling network to the sex-chromosome architecture. These findings provide a two-pathway framework for understanding the hormonal basis of sexual plasticity in *C. sativa* and identify specific GA-related genes as candidates for functional validation and potential targets for breeding sex-stable cultivars.

## Acknowledgments

The authors gratefully acknowledge the support of the Natural Sciences and Engineering Research Council (NSERC) Discover Grant number RGPIN-2022-03396. JR has been supported by the Fonds de recherche du Québec (https://doi.org/10.69777/2009140). ASM has also been supported by a NSERC Canada Vanier Graduate Scholarship.

## Competing Interests

The authors have no competing interests to declare.

## Author Contributions

Conceptualization: ASM and JR

Data Curation: ASM and JR

Formal Analysis: ASM and JR

Funding Acquisition: DT

Investigation: JR

Methodology: ASM and JR

Project Administration: ASM and DT

Resources: DT

Software: JR

Supervision: DT

Visualization: JR

Writing-Original Draft Preparation: ASM and JR

Writing-Review and Editing: ASM, JR and DT

## Data Availability Statement

All WGS and RNA-seq data analyzed in this study have been deposited in the NCBI Sequence Read Archive (SRA) and are publicly available under BioProject accession PRJNA1404156. All scripts and raw data used in this study can be found under the Open Science Framework (OSF) project https://doi-org.acces.bibl.ulaval.ca/10.17605/OSF.IO/X5CNY.

## Notes

https://www.ncbi.nlm.nih.gov/sites/myncbi/julien.roy.2/collections/63974920/public/

### Competing Interest Statement

The authors have declared no competing interest.

